# Deciphering ESR1-driven transcription in human endometrial stromal cells via transcriptome, cistrome, and integration with chromatin landscape

**DOI:** 10.1101/2025.04.23.650343

**Authors:** Skylar G. Montague Redecke, Austin Bell-Hensley, Shuyun Li, MyeongJin Yi, Akshadha Jain, Abdull J. Massri, Francesco J. DeMayo

**Author notes:** **Corresponding author:** Francesco J. DeMayo, Reproductive and Developmental Biology Laboratory, National Institute of Environmental Health Sciences, NIH, Durham, NC, USA. Mailing Address: 111 TW Alexander Dr Bldg: David P Rall Building Rm: B304 Research Triangle Park, NC 27709. **Disclosure Statement:** Austin Bell-Hensley: Nothing to disclose. Shuyun Li: Nothing to disclose. MyeongJin Yi: Nothing to disclose. Akshadha Jain: Nothing to disclose. Abdull J. Massri: Nothing to disclose. **CRediT Authorship Contribution Statement:** Austin Bell-Hensley: Formal analysis, Visualization. Shuyun Li: Writing – Review & Editing. MyeongJin Yi: Investigation. Akshadha Jain: Investigation. Abdull J. Massri: Formal analysis. **Attestation Statement:** Data regarding any of the subjects in the study has not been previously published unless specified. Data will be made available to the editors of the journal for review or query upon request pre and/or post publication.

## Abstract

**Objective:** To investigate ESR1 and estrogen-driven transcription in human endometrial stromal cells.

**Design:** Telomerase-immortalized human endometrial stromal cells were engineered to activate ESR1 using the CRISPR activation system and treated with vehicle or E2. Primary endometrial stromal cells were treated with vehicle or decidualization cocktail.

**Subjects:** Biopsies from two healthy, reproductive-aged volunteers with regular menstrual cycles and no history of gynecological malignancies.

**Main Outcome Measures:** Bulk RNA-sequencing in ESR1-activated and E2-treated cells was compared to identify ligand-independent and -dependent ESR1 actions. Cut&Run was performed in ESR1-activated cells treated with E2 or vehicle to assess the ESR1 cistrome. H3K27ac HiChIP was conducted in primary endometrial stromal cells treated with vehicle or decidualization cocktail to identify hormonally induced chromatin interaction changes.

**Results:** Among six tested guide RNAs, the ESR1-3 gRNA induced robust ESR1 activation, which restored E2 responsiveness in THESC. Bulk RNA-seq revealed ESR1-mediated E2-dependent and E2-independent transcriptional programs, regulating pathways involved in inflammation, proliferation, extracellular matrix organization, and cancer. Notably, 72% of differentially expressed genes (DEGs) overlapped with genes active in human endometrial tissue during the proliferative E2 dominant phase, supporting their physiological relevance. Cut&Run-seq identified genome-wide ESR1 binding sites, with most located at distal regulatory elements. To associate distal ESR1 binding sites with genes, we integrated H3K27ac HiChIP chromatin loops in hESC to identify distal ESR1 binding sites that loop to gene promoters. We identified genes regulated by ESR1/E2 through long-range chromatin looping that are involved in stromal cell decidualization, including FOXO1 and IL6R. Additionally, we identified genes implicated in endometrial cancer, including ERRFI1, NRIP1, and EPAS1, suggesting a role for stromal ESR1 driven endometrial pathologies. Functional assays confirmed that ESR1 promotes cell viability and, in the presence of E2, enhances migration.

**Conclusions:** ESR1 activation through CRISPR restores E2 responsiveness in endometrial stromal cells. Changes to chromatin architecture support gene expression changes that drive decidualization. Integration of ESR1/E2 transcriptome and cistrome with HiChIP data identifies its role in regulating inflammation, proliferation, and decidualization, as well as its implications in endometrial cancer. This model serves as a powerful tool to study estrogen signaling in endometrial stromal biology and related pathologies.

## INTRODUCTION

The endometrium consists of a luminal epithelial layer overlying a connective tissue matrix primarily comprised of endometrial stromal cells, with populations of endothelial cells and immune cells (1). It undergoes dynamic changes throughout the menstrual cycle in response to sex steroid hormones estrogen and progesterone, preparing for potential embryo implantation and supporting early pregnancy (2).

Estrogen, specifically estradiol (E2), is a key regulator of endometrial function, promoting endometrial proliferation after menstruation and preparing the endometrium to become progesterone responsive for implantation and decidualization (3). The menstrual cycle is regulated by the tightly coordinated release of estrogen and progesterone in two main phases: the proliferative and the secretory phase. During the proliferative phase, E2 signaling from the stroma promotes the thickening of the endometrium by triggering proliferation in both stromal cells as well as epithelial cells in a paracrine manner (4, 5). E2 also activates the progesterone receptor gene (PGR), preparing the endometrium to respond to progesterone (P4) secreted by the corpus luteum during the secretory phase (6). A rise in P4 signaling during the secretory phase, alongside E2 signaling, creates a receptive endometrium initiating a process known as decidualization, preparing it for implantation (7). During decidualization, fibroblast-like endometrial stromal cells differentiate into epithelioid-like decidual cells that create an immunotolerant environment and provide nutritional support for the implanting blastocyst prior to placentation (8). While P4 is the primary hormonal regulator of decidualization, E2 signaling is also crucial for its success, but the direct mechanisms through which E2 regulates this process remain unclear (9).

E2 primarily exerts its effects in the female reproductive tract by binding to nuclear estrogen receptors, ERα and ERβ, which are encoded by the ESR1 and ESR2 genes, respectively. Both ESR1 and ESR2 are expressed in the ovaries, the endometrial stroma, and the epithelium, but ESR1 is the dominant isoform throughout the endometrium (10). Knockout of endometrial-specific ESR1 in mice leads to infertility due to defects in implantation and decidualization (11–14). ESR1 can regulate gene transcription by directly binding to genomic estrogen response elements (EREs) or by interacting with other transcription factors bound to DNA regions lacking EREs (13).

Dysregulated E2 signaling is associated with impaired fertility and reproductive diseases including type I endometrial tumors (3), endometriosis, and polycystic ovarian syndrome (15–17). Endometrial cancer is the most common malignancy of the female reproductive tract and primarily affects post-menopausal women, a period characterized by unopposed estrogen signaling due to a decline in progesterone levels (American Cancer Society, endometrial cancer, https://www.cancer.org). Approximately 75% of endometrial tumors are type 1, characterized by overactivation of ESR1 expression, and are hypothesized to be estrogen-driven (3). Therefore, understanding the E2/ESR1 signaling is crucial for successful pregnancy and developing targeted therapies for endometrial diseases.

While transcriptomic and cistromic studies of ESR1 have been extensively conducted in mice and humans using whole endometrial biopsies, the presence of both epithelial and stromal components in these samples makes it challenging to isolate the specific role of ESR1 in the stroma. Endometrial epithelial cells and organoid models have been widely used to study E2 signaling in the epithelium in vitro, as they exhibit a strong response to E2 (18). However, studying ESR1 function in the endometrial stroma remains challenging due to the low expression of ESR1 in primary and immortalized human endometrial stromal cells (hESC), which limits E2 responsiveness (19). To overcome this barrier, we engineered telomerase-immortalized hESCs (THESCs) with a CRISPR activation system to restore ESR1 expression, re-establishing E2 responsiveness in hESCs in vitro. We investigated the ligand dependent and independent ESR1 transcriptome and cistrome through bulk RNA-sequencing (RNA-seq) and Cleavage Under Targets and Release Using Nuclease (Cut&Run) assays. We also examined the chromatin landscape in hESC using H3K27ac HiChIP and demonstrated the utility of this dataset by integrating it with RNA-seq and Cut&Run to identify E2 target genes with distal ESR1 binding that loop to gene promoters.

Functionally, ESR1 activation promoted cell viability and migration in response to ESR1/E2, activating genes associated to endometrial cancer, including ERRFI1, NRIP1, and EPAS1. These findings reveal the ESR1-driven transcription in hESC, providing critical insights into fertility and endometrial pathologies.

## MATERIALS AND METHODS

### Cell culture and engineering

Telomerase-transformed human endometrial stromal cells, THESC (20) (ATCC, CRL-4003), were maintained in DMEM/F-12 (Gibco Thermo Fisher Scientific, Waltham, MA, USA) supplemented with 10% Fetal Bovine Serum (FBS, Gibco Thermo Fisher Scientific) and antibiotics (10 000 IU/mL penicillin, 10 000 IU/ mL streptomycin; Life Technologies, Grand Island, NY, USA), termed regular hESC media. Telomerase transformed human myometrial cells (hTERT-HM) and stromal cells (H1644) were maintained according to their respective protocols (21). Cell culture media was filtered using the 0.22um Rapid-Flow™ Sterile Disposable Filter Units (Nalgene, Thermo Fisher Scientific). Cells were cultured and grown in a 5% CO_2_ and 37°C incubator. Cell media was changed every three days and cells were passaged before reaching 90% confluency.

To activate ESR1, we used a split-plasmid CRISPR activation (CRISPRa) system which utilizes two separate plasmids that are introduced into cells, one carrying a guide RNA (gRNA) and the other carrying a dead Cas9 enzyme fused to the transcriptional activators VP64, p65, and Rta (dCas9-VPR). To introduce the dCas9-VPR and gRNA plasmids into uterine cells, we used lentivirus transduction. Ef1a-dCas9-VPR-Blast was obtained as prepackaged lentivirus from Dharmacon (CRISPRmod CRISPRa lentiviral dCas9-VPR, Dharmacon/Horizon Discovery, Lafayette, CO, USA). Cells were engineered following the manufacturers protocol with the following specifications.

50,000 cells per well were seeded in 24-well plates and cultured in regular hESC media. 24 hours later, media was changed to 0.25mL of DMEM/F12 without antibiotics or serum and transduced with lentiviral particles at a multiplicity of infection (MOI) of 4.0. After 4-6 hours of incubation, 0.75mL regular hESC media was added to each well and cells were incubated for 48 hours. Subsequently, cells were cultured in DEMEM/F-12 (Gibco Thermo Fisher Scientific) containing 10% FBS (Gibco Thermo Fisher Scientific) and 4 ug/mL Blasticidin (Gibco R21001, Thermo Fisher Scientific), termed engineered hESC media, for 6-8 days to select for cells expressing dCas9-VPR; cells were passaged as necessary. Cells were grown up in a T175 and frozen down in aliquots. For experiments, THESC^dCas9-VPR^ were thawed and incubated for at least two days prior to transduction with gRNA, and cells were used for experiments within 10 passages from thawing frozen stock.

### gRNA design, lentivirus production, and cell transduction

gRNAs were designed using the CHOPCHOP (22, 23) and CRISPick (24) tools to target upstream of alternate transcription start sites annotated in the NCBI Refseq database (Table 1.1). All gRNA expression vectors were synthesized by and acquired from VectorBuilder (VectorBuilder.com) and expressed neomycin resistance and GFP markers. gRNA expressing plasmids were isolated from bacteria using Plasmid Midi Kit (Qiagen, Hilden, Germany) according to the manufacturer’s instructions. Lentivirus was generated by transfection of the constructs and packaging vector into HEK293T cells with the support of the NIEHS Viral Vector Core Facility. For viral transduction, cells were seeded at 20% confluency in 10 cm plates, 24 hours before transduction. Cells were transduced with gRNA lentivirus at an MOI of 12 and incubated for 24 hours before washing plates with PBS and replacing with engineered hESC media.

**Table 1.1:**
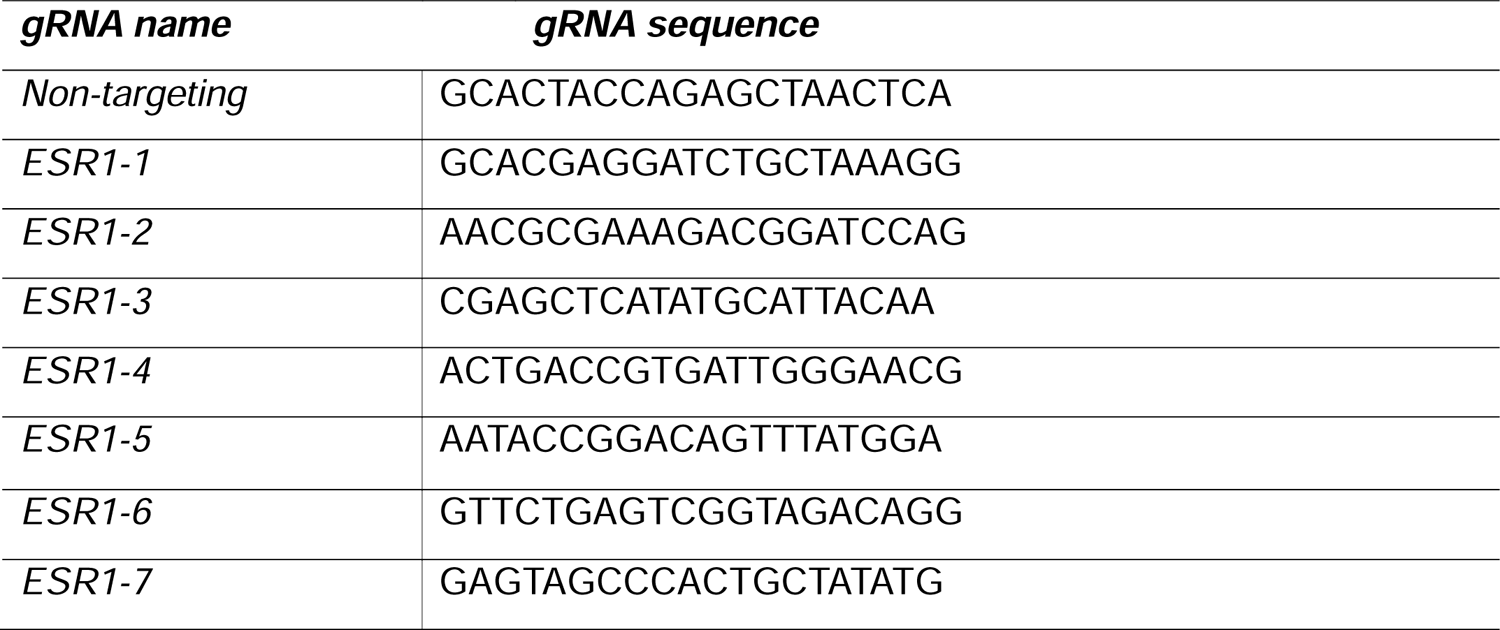
gRNA sequences.

THESC^ESR1^ were maintained in engineered hESC media and used for experiments within three weeks of transduction. gRNA expression was assessed by GFP expression.

### 17β-Estradiol treatment

Wells/plates were washed with PBS and media was replaced with 3 mL OptiMEM (Gibco Thermo Fisher Scientific) with 2% Charcoal Stripped Serum (csFBS, Gibco Thermo Fisher Scientific) and 1% Penicillin-Streptomycin), termed low serum media. Low serum media was supplemented with either 10nM 17β-estradiol (Sigma-Aldrich, St. Louis, MO, USA), termed E2 media, or 0.01% Ethanol, termed vehicle media. Cells were collected at different time point (indicated under each protocol) for isolating RNA (RT-qPCR and RNA-seq), or conducting Cut&Run, Transwell Migration Assay, MTT assay, or colony formation assay.

### Western blot

Protein was isolated and quantified as cited in Wu et al., {Wu, 2025 #99} using RIPA Lysis and Extraction Buffer (#89900, Thermo Fisher Scientific). 50ug of protein was separated on 4-20% Criterion TGX Precast Gels (# 5678094, BioRad, Hercules, CA, USA) with the Precision Plus Protein Dual Color Standards Ladder (#1610374, BioRad). Gels were transferred to nitrocellulose membrane using the Turbo-Blot transfer system (BioRad) according to the manufacturer’s protocols. Membranes were blocked as cited in Wu et al., {Wu, 2025 #99}. Antibodies to ESR1 (Cell Signaling Technologies (D6R2W), #13258, Danvers, MA, USA; 1:500) or GAPDH (Santa Cruz Biotech, sc-25778, Dallas, TX, USA; 1:1000) were diluted in blocking buffer and incubated with membranes overnight at 4°C. After 3 5-minute washes with TBST, anti-Rabbit IgG or anti-Mouse IgG labeled with IR 800 (LI-COR Biosciences, Lincoln, NE, USA) was diluted 1:20,000 in blocking buffer and incubated with membranes for 45 minutes at room temperature. After 3 5-minute washes with TBST, membranes were imaged using Odyssey Fc Imager (LI-COR Biosciences) using 800nm channel for 10 minutes, and 700nm channel for 30 seconds. Bands were quantified using Image Studio v5.5.

### RNA isolation and qPCR

The total RNA of the cells was isolated using Qiagen RNeasy RNA mini prep kit with Qiagen on-column DNase digestion following the manufacturer’s instructions. 1 µg of total RNA was reverse transcribed into cDNA using Moloney Murine Leukemia Virus reverse transcriptase (Thermo Fisher Scientific) with Random Hexamers (Invitrogen, Waltham, MA, USA) according to manufacturer protocol. Quantitative real time PCR was performed using SsoAdvanced Universal SYBR Green Supermix (BioRad). Briefly, reaction samples were prepared to a total volume of 20ul with 5uM of each of the forward and reverse primers, 4ul of cDNA, and a final 1x concentration of the SYBR Green Supermix. SYBR green primers were obtained from PrimerBank (https://pga.mgh.harvard.edu/primerbank/) or designed using NIH Primer-blast (25) and synthesized by Sigma-Aldrich (Table 1.2). The reaction was heated to 95°C for 30 seconds, followed by 40 cycles of denaturation at 95°C for 15 seconds and annealing and elongation at 60°C for 30 seconds. Temperature cycles were performed on the CFX Connect^TM^ Real-Time PCR Detection System (BioRad). Each reaction was performed in technical duplicates and ΔΔCt values were calculated using 18S control amplification results to determine the relative mRNA levels per sample.

### Transwell migration assay

Cells were seeded in 10cm plates and cultured in engineered hESC media until reaching 60% confluence. Plates were washed with PBS and media was changed to E2 media or vehicle media. 24 hours later, 30,000 cells in 300uL of E2 or vehicle media were seeded in 8.0 um transparent PET membranes (Corning, Falcon, 353097, Corning, NY, USA) placed in 24-well plates. 700uL of regular hESC media was added to the bottom of the plates. After 48 hours of culture, the cells on the top side of the membrane were wiped with a cotton swab dipped in PBS. The insert was fixed with 4% paraformadehyde in PBS for 15 minutes, washed with PBS two times for 5 minutes each, stained with 1% crystal violet (Sigma-Aldrich) for 20 minutes, then washed with PBS three times for 10 minutes each before imaging on a brightfield microscope. Intensity of staining was quantified using ImageJ.

### MTT assay

1,000 cells were seeded in 96-well plates and cultured in engineered hESC media for 24 hours. Wells were washed with PBS and media was changed to 100 uL per well E2 media or vehicle media. Media was changed every two days and cells were incubated for a total of 4 days. Viability was assessed with a Cell Proliferation Kit I (MTT) (Roche, Basel, Switzerland) according to the manufacturer’s protocol. Briefly, 10 uL of the MTT labeling reagent (final concentration 0.5 mg/mL) was added per well and incubated for 4 hours before adding 100 uL of the Solubilization buffer and incubating overnight.

Absorbance at 570 nm was measured and relative cell viability was calculated by subtracting absorbance at 570 from control wells containing media only with no cells and subtracting reference wavelength at 670nm.

### RNAseq and data analysis

100,000 THESC^ESR1^ cells per well were seeded in 6-well plates and cultured in engineered hESC media for 24 hours. Wells were washed with PBS and media was changed to 3 mL per well E2 media or vehicle media for 24 hours. The total RNA of the cells was isolated using Qiagen RNeasy RNA mini prep kit with Qiagen on-column DNase digestion following the manufacturer’s instructions. Libraries were prepared and sequenced as 75 bp paired-end reads by the NIEHS Sequencing Core using the Illumina Tru-seq Stranded mRNA library kit and Illumina NovaSeq 6000 instrument.

For RNA-seq analysis, adapter trimming, optical deduplication, and PCR deduplication were performed using BBMap v39.01 and the trimmed reads were subsequently mapped to the hg38 genome (GCF_000001405.40) using STAR v2.6.0c. Genes with fewer than 200 reads across all samples were excluded, and differential expression analysis was performed using edgeR v4.2.2 in R v4.4.1. Functional analysis of each gene list was performed using Ingenuity Pathway Analysis software v01.22.01 (IPA, www.ingenuity.com) and Metascape version X0200 (http://metascape.org).

For previously published RNA-seq (GSE205481) (26), raw Fastq files were downloaded from GEO (samples P1, P2, and P3 siNT-Pre-dec and siNT-Dec) and analyzed following the same protocol as aforementioned. For previously published RNA-seq (GSE132713) (27), normalized counts were downloaded from GEO and filtered to include genes with average FPKM > 1 in the human endometrium proliferative samples.

### Cut&Run and data analysis

The ESR1 Cut&Run samples were prepared with the CUTANA^™^ ChIC/CUT&RUN Kit (Version 3.5; EpiCypher, Durham, NC, USA). Briefly, cells were treated with E2 media or vehicle media for 1, 3, or 6 hours. A total of 550,000 cells per sample were collected in DNA LoBind 1.5mL tubes (Eppendorf, Hamburg, Germany) containing wash buffer.

Cells were captured with BioMag Plus Concavalin A beads (Bangs Laboratories, Inc, Fishers, Indiana, USA) and incubated with ESR1 antibody (Cell Signaling Technologies, #13258, Danvers, MA, USA; 1:25) or IgG antibody (1:100) supplied by the CUTANA kit overnight at 4°C in antibody incubation buffer. After washing away antibodies, pAG-MNase generated by the NIEHS proteomics core was added and incubated for 30 minutes at room temperature. After washing, CaCl2 was added and incubated for 30 minutes in an aluminum cold block placed in ice. The protein-DNA complexes were released by incubating at 37°C for 10 minutes. DNA was purified using the Qiagen MiniElute PCR Purification Kit. Sequencing libraries were prepared using the NEBNext Ultra II DNA library preparation kit for Illumina (New England Biolabs, Ipswich, MA, USA) according to the manufacturer’s instructions, with 14 cycles of PCR amplification. The PCR products were quantified using the Qubit dsDNA HS Assay Kit (Invitrogen, Waltham, MA, USA) and quality control was performed with Bioanalyzer DNA Analysis (Agilent Technologies, Santa Clara, CA, USA). The libraries were sequenced by the NIEHS sequencing core.

Reads from CUT&RUN were trimmed of adapter sequences using Trim Galore! (https://www.bioinformatics.babraham.ac.uk/projects/trim_galore/; Version 0.6.10) with default settings except for specifying Illumina adapter sequences and paired-end validation. Trimmed sequences were aligned to hg38 and E. coli K-12 MG1655 genomes using bowtie2 (Version 2.5.2) with maximum fragment length 2,000, forward-reverse mate orientation, local read alignment, and no mixed or discordant read alignments (28). Alignment files were processed using Picard (http://broadinstitute.github.io/picard; Version 3.2.0) for duplicate removal and samtools (Version 1.18) to filter for reads with MAPQ ≥ 5 (29). HOMER v5.1 was used to call peaks for each sample with default parameters except mode of operation set to factor and single-end reads. Peak calls for all samples were collapsed to a set of union peaks which were subsequently assigned as positive or negative for ESR1 signal per sample at a threshold of 5x the median IgG signal (30). Only peaks present in both replicates of a condition were used for downstream analysis. Peaks were annotated for overlaps with HiChIP loop anchors and genomic features in the NCBI RefSeq gene model as of February 06, 2025 (31). In cases where a peak overlaps multiple genomic features, only the highest priority genomic feature was annotated using the following priority rankings: promoter, 5’ UTR, 3’ UTR, exon, intron, intergenic. To identify overlapping and unique peaks between samples, we used the HOMER mergePeaks function with a maximum merge distance of 100 nucleotides. For Motif enrichment analysis, we used the HOMER findMotifsGenome.pI function, using the “-size given -p 10” parameters. To identify genes regulated by ESR1/E2 with nearby ESR1 binding, genes within 100 kb of a peak (upstream or downstream) were overlapped with E2-dependent DEGs for pathway analysis.

### HiChIP and data analysis

Primary endometrial stroma cells (hESCs) were isolated from endometrial biopsies collected under the human subject protocol number H-13062 approved by the institutional review board of Baylor College of Medicine. The biopsies were obtained from two healthy, reproductive-aged volunteers with regular menstrual cycles and no history of gynecological malignancies. All donors provided written informed consent. hESCs were maintained as cited in Li et al., (2023). To induce decidualization, hESC were treated with 0.2% ethanol (vehicle) or a mixture of 10 nM 17β-estradiol (Sigma-Aldrich), 1 µM medroxyprogesterone acetate (Sigma-Aldrich) and 100 µM dibutyral cyclic AMP (Sigma-Aldrich), in low serum media, termed EPC, for 3 days. HiChIP libraries were prepared by Arima Genomics and sequenced by the NIEHS sequencing core.

The HiCUP platform (Version 0.9.2) was used to process and align HiChIP samples with Arima digest and default parameters except minimum and maximum ditag length (set to 100 and 1,000 bp) (32). Peaks were inferred from the HiChIP data using MACS2 (Version 2.2.9.1) with a q-value threshold of 0.01 and extension size of 100 and the -- nomodel and --keep-dup all flags (33). Loops with at least one loop anchor in a peak region were called and differential loops were identified using FitHiChIP (Version 11.0) following the author’s vignette except for bin size (set to 10 kb) and background model (set to peak-to-all; option 0) (34). Loop anchors were annotated for overlapping genes and genomic features in the NCBI RefSeq gene model as of February 06, 2025 (31). In cases where loop anchors overlap multiple genomic features, only the highest priority gene and genomic feature were annotated for each loop anchor using the following priority rankings: promoter, 5’ UTR, 3’ UTR, exon, intron, intergenic. These annotations were used to identify ESR1 peaks that were one loop away from a promoter. Pathway analysis was performed on E2-dependent DEGs whose promoters looped to an ESR1 binding site. HiChIP visualizations were generated with Juicer and Juicebox for interaction matrices and R package Sushi for loop plots (35–37).

### Statistical Analysis

Obtained data were expressed in a mean ± standard deviation of the 2-5 biological replicates. Statistical analysis was performed using GraphPad Prism (Version 10.4.0 (621), GraphPad Software Inc.). Distribution of datapoints was assessed by Shapiro-Wilk test. If datapoints were normally distributed, statistics were performed by One-Way ANOVA Turkey test for more than two comparisons, and parametric unpaired t test for two comparisons. Significance was defined as p-value < 0.05.

## RESULTS

### Activating ESR1 in THESC using the CRISPR activation system

To overcome the limited responsiveness of THESC to E2, we activated ESR1 using a dCas9-VPR plasmid with a blasticidin resistance gene (Figure 1A). RT-qPCR confirmed successful transduction and expression of dCas9 in comparison to untransduced THESC; these cells will be referred to as THESC^dCas9-VPR^ from this point forward (Figure 1B). Due to a reduced expression of dCas9 after 8 days without selection, we maintained THESC^dCas9-VPR^ in media always containing blasticidin to ensure sustained expression (Figure 1B). To activate ESR1, we designed seven different gRNAs (ESR1-1 to ESR1-7) to target alternate ESR1 transcription start site (TSS) annotated in the NCBI Refseq database (Figure 1C). RT-qPCR and western blot showed that the ESR1-3 gRNA can successfully activate ESR1 in comparison to the non-targeting control gRNA (NT gRNA) (Figure 1C, D, S1). These cells will be referred to as THESC^ESR1^ and THESC^NT^ from this point forward. We also show that dCas9-VPR and ESR1-3 gRNA can successfully activate ESR1 in myometrial cells (hTERT-HM) and a second endometrial stromal cell-line (H1644) (Figure S2A-B). To assess ESR1 functionality, we treated THESC^ESR1^ and THESC^NT^ with 10 nM E2 for 24 hours and tested expression of the known estrogen target gene PGR, as well as the ESR1 target gene INPP4B (19). By RT-qPCR, we observed an increase of PGR mRNA expression in response to E2, and an increase in INPP4B mRNA expression following ESR1 activation (Figure 1E). These results confirm that ESR1 can successfully be activated using the ESR1-3 gRNA and that the cells are ESR1/E2 responsive.

**Figure 1.**
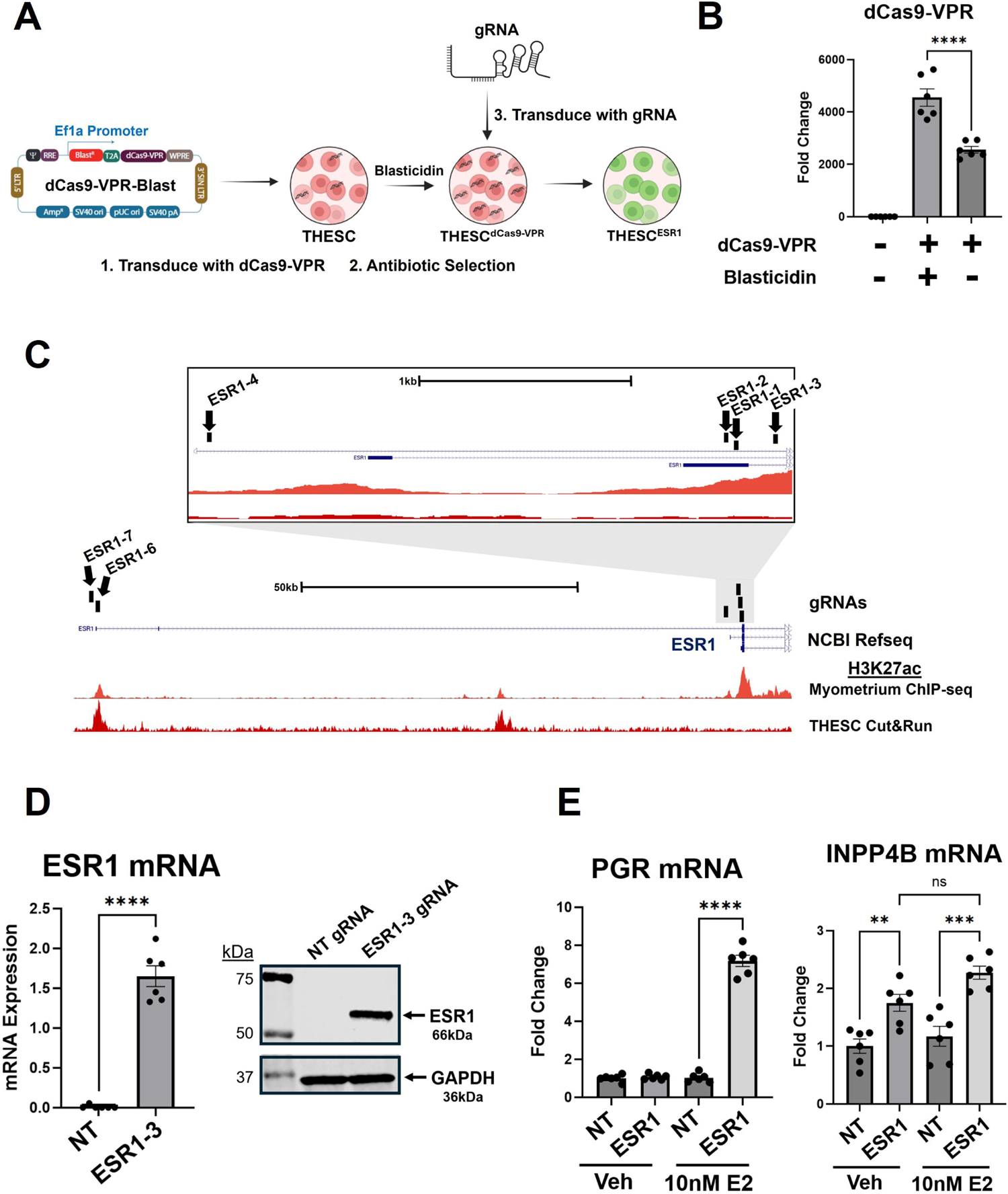
Activating ESR1 in immortalized human endometrial stromal cells (THESC) using CRISPRa. (A) Schematic for engineering cells with split plasmid CRISPR activation system using dCas9-VPR-Blast. (B) RT-qPCR analysis of dCas9-VPR expression in non-transduced cells, dCas9-VPR transduced cells cultured in blasticidin for 8 days, and dCas9-VPR transduced cells cultured in the absence of blasticidin for 8 days. Error bars are shown as mean with SEM. (C) UCSC genome browser view showing the location of the six gRNAs designed to target near alternate ESR1 transcription start sites annotated in the NCBI Refseq database. Tracks show H3K27ac ChIP-seq in myometrial tissue (orange) and H3K27ac Cut&Run in THESC (red) (unpublished data). (D) RT-qPCR analysis of ESR1 expression in THESC^dCas9-VPR^ transduced with ESR1-3 gRNA (THESC^ESR1^) in comparison to control NT gRNA (THESC^NT^). (E) Western blot analysis of ESR1 and GAPDH expression in THESC^ESR1^ and THESC^NT^. (G) RT-qPCR analysis of ESR1 target genes in THESC^ESR1^ and THESC^NT^ treated with vehicle or 10 nM E2 for 24 hours. Statistical significance was determined by student’s t-test for two comparisons, and one-way ANOVA for more than two comparisons, with significance defined as p-value < 0.05. THESC = immortalized human endometrial stromal cells; SEM = standard error of the mean; TSS = transcription start site; THESC = immortalized human endometrial stromal cells; E2 = 10 nM estradiol.

### ESR1 E2-dependent and E2-independent transcriptomic regulation in engineered THESC

To identify ESR1 E2-dependent and -independent target genes, THESC^ESR1^ and THESC^NT^ were treated with 10 nM E2 or 0.01% ethanol (vehicle) for 24 hours and bulk RNA-seq was conducted to observe changes in gene expression. Hierarchical Clustering and Multidimensional Scaling (MDS) plot reveal distinct clustering of THESC^ESR1^ and THESC^NT^ in the presence and absence of E2, suggesting that ESR1 regulates gene expression in ligand dependent and independent manners, and that ESR1 activation restores E2 responsiveness (Figure S3A-B).

ESR1 transcriptomic regulation was separated into E2-independent and E2-dependent ESR1 and DEGs were identified as genes with absolute fold-change (FC) > 1.3 and adjusted p-value < 0.05 (Figure 2A). A total of 305 E2-independent DEGs were identified and ESR1 was the top activated gene (FC > 637) (Figure 2B; Dataset 1). IPA pathway analysis indicated an enrichment of pathways related to cancer, inflammation, proliferation, migration, angiogenesis, preeclampsia, and extracellular matrix organization (Figure 2C; Dataset 1). A total of 369 E2-dependent DEGs were identified (Figure 2D; Dataset 2), and the long non-coding RNA LINC01016 was the top activated gene (FC > 37; not shown in figure because out of scale). This gene is associated with endometrial cancer (38) and breast cancer and is a direct target of ESR1 in MCF7 and T47D breast cancer cells (39). IPA revealed enrichment of pathways related to cancer, inflammation, axonal guidance, metabolism, wound healing, and Wnt/β-catenin signaling (Figure 2E; Dataset 2). To evaluate the relevance of these RNA-seq datasets in relation to in vivo human tissue, DEG lists were compared to active genes (FPKM > 1) in human endometrial tissue during the proliferative stage (GSE132713), the stage when estrogen signaling is dominant (27). Overall, 72.9% and 72.1% of the E2-dependent and independent DEGs, respectively, were active in the human endometrium during the proliferative stage, supporting the significance of the genes identified using engineered THESC (Figures 2F; Dataset 3). All together, these results suggest that ESR1 regulates gene expression in both the presence and absence of E2, and triggers gene expression changes involved in endometrial stromal cell biology and function.

**Figure 2.**
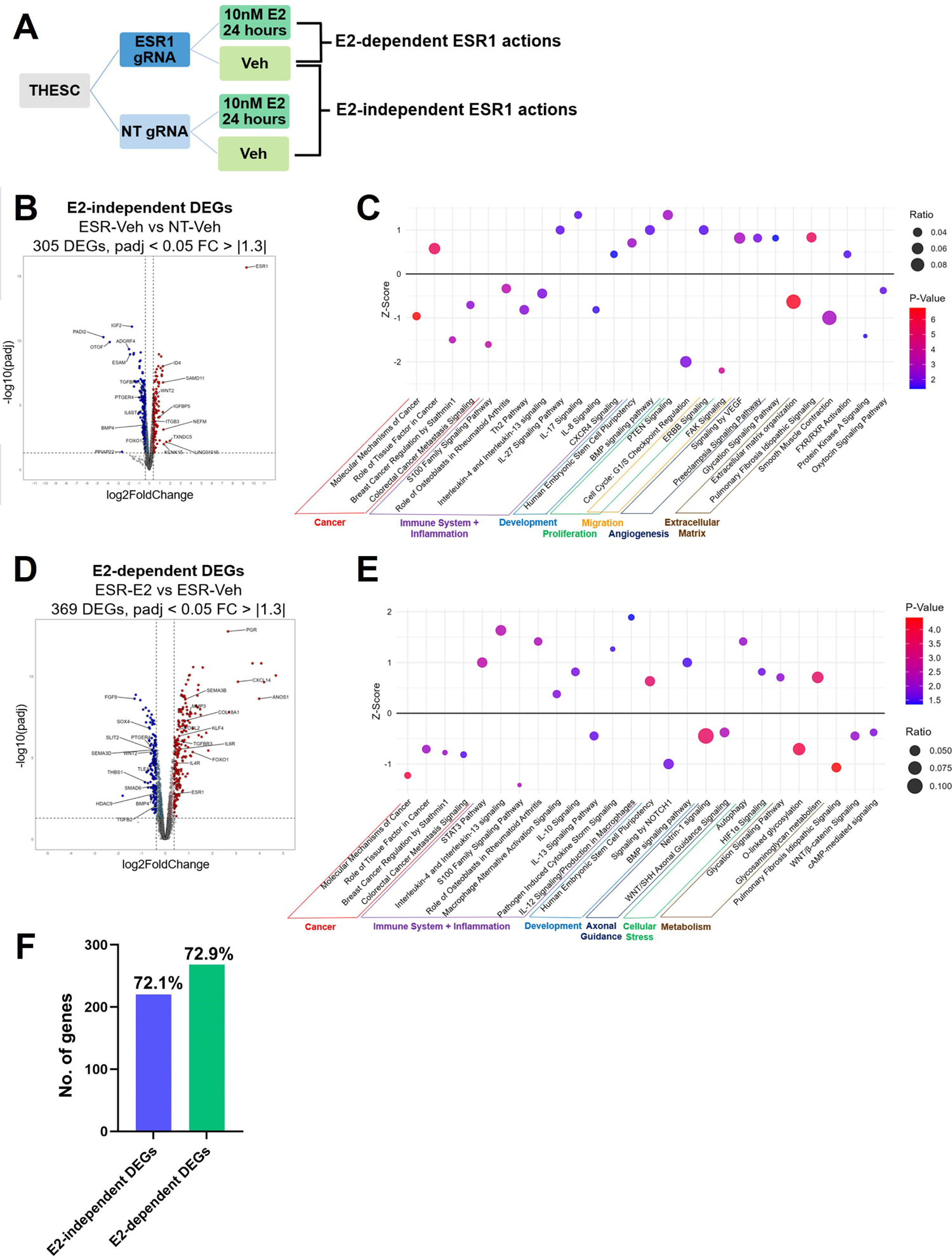
ESR1 E2-dependent and E2-independent transcriptomic regulation in engineered THESC. (A) Schematic of comparisons to investigate E2-dependent and independent actions in THESC by bulk RNA-seq. (B) Volcano plot representation and (C) IPA pathway analysis of differentially expressed genes in response to unliganded ESR1 by comparing vehicle-treated THESC^ESR1^ and THESC^NT^. (D) Volcano plot representation and (E) select pathways from IPA pathway analysis of differentially expressed genes in response to liganded ESR1 by comparing THESC^ESR1^ treated with 10 nM E2 or vehicle. (F) Bar graph showing the number and percentage of E2-independent and dependent DEGs that are active (FPKM > 1) in human endometrial biopsies at the proliferative stage from published RNA-seq data (GSE132713). THESC = immortalized human endometrial stromal cells; THESC^ESR1^ = ESR1 activated cells; THESC^NT^ = control non-targeting cells; E2 = 10 nM estradiol; vehicle = 0.1% ethanol; IPA = ingenuity pathway analysis

### ESR1 E2-dependent and E2-independent cistrome in engineered THESC

To investigate the ESR1 cistrome, we performed ESR1 Cut&Run in THESC^ESR1^ treated with 10 nM E2 or vehicle for 1, 3, or 6 hours (Figure 3A). ESR1 had the greatest number of binding peaks at the 3-hour timepoint (1305), encompassing nearly all the peaks from the 1-hour and 6-hour E2 timepoints (Figure 3B). At 3-hours, 97% of ESR1 peaks without E2 were also present with E2 (Figure 3C), consistent with previous findings showing minimal positional differences in ESR1 binding in the presence and absence of E2 in the mouse uterus (40). Thus, we decided to focus on ESR1 peaks after treatment with E2 for 3-hours. Most peaks (54.1%) were located distal to genes (>25 kb from the TSS) and within intergenic regions (Figure 3D), consistent with previous studies showing that ESR1 binds primarily to distal cis-regulatory elements (41). 58% of ESR1 peaks in THESC overlapped with ESR1 ChIP-seq peaks from proliferative-stage human endometrium (GSE200807) (18) (Figure 3E). Differences may reflect paracrine signaling between stromal and epithelial cells in biopsies that are absent in monoculture. Motif enrichment analysis using HOMER identified estrogen response element (ERE), the canonical ESR1 binding motif, as the most enriched motif, along with motifs that bind Jun-AP1, RUNX, and TEAD (Figure 3F) which have been shown to mediate ESR1 genomic binding and transcriptional regulation in breast cancer cells (42–44). These results support previous findings that ESR1 binds to distal cis-regulatory elements and regulates gene expression through a tethering mechanism via protein-protein interactions (45).

**Figure 3.**
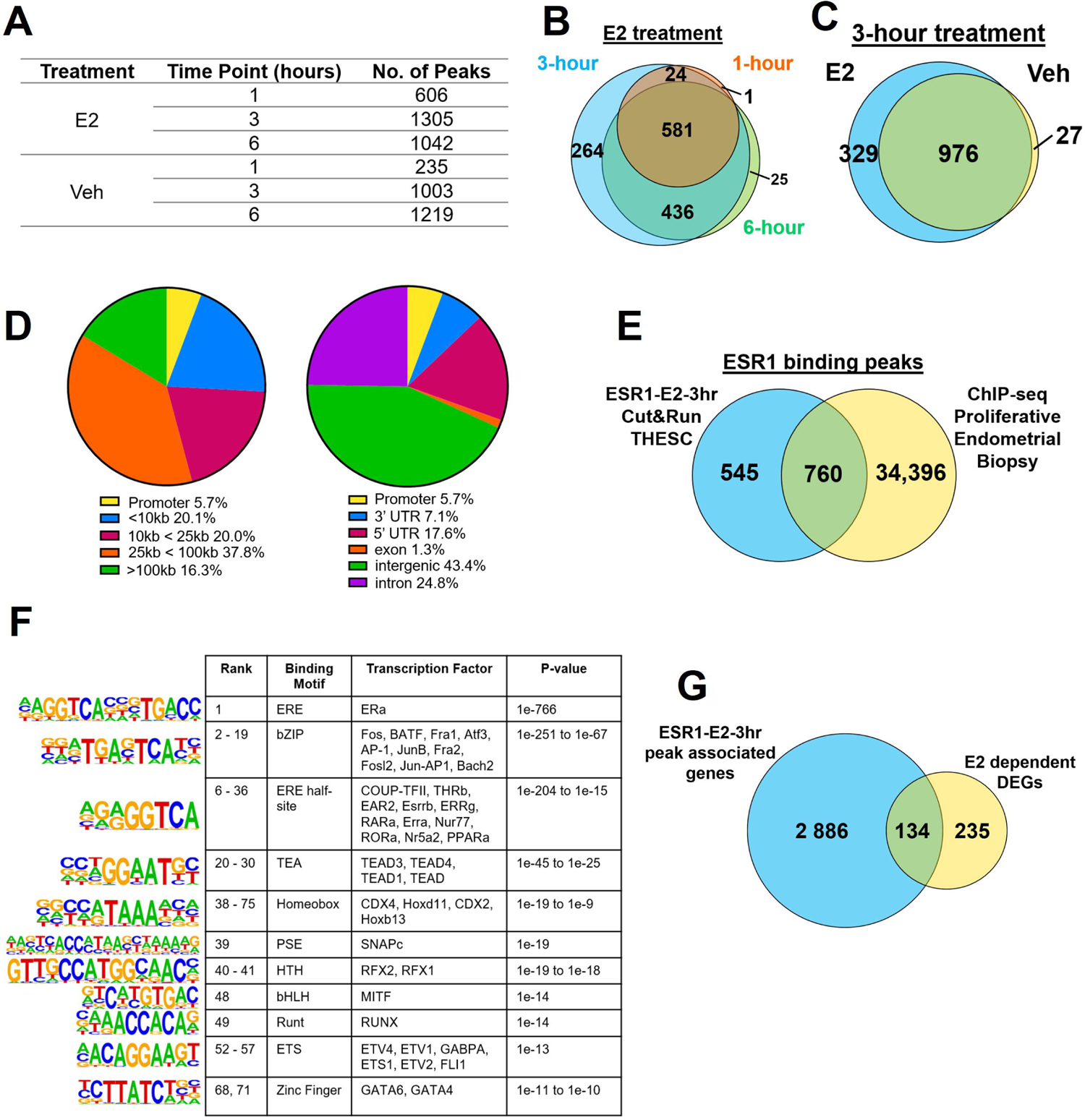
ESR1 E2-dependent and E2-independent cistrome in engineered THESC. (A) Table showing the number of ESR1 peaks in THESC^ESR1^ after treatment with E2 or vehicle for 1, 3, or 6 hours. Experiment was performed in biological duplicates and only peaks present in both duplicates were reported. (B) Venn diagram comparing number and location of ESR1 peaks after treatment with E2 for 1, 3, or 6 hours. (C) Venn diagram comparing number and location of ESR1 peaks in the presence or absence of E2 at the 3-hour timepoint. (D) Distance of ESR1 peaks (treated with E2 for 3 hours) from the nearest TSS, and location relative to annotated genes. (E) Venn diagram comparing number and location of ESR1 peaks in THESC^ESR1^ (treated with E2 for 3 hours) and ESR1 ChIP-seq peaks in proliferative endometrial biopsies from previously published data (GSE200807). (F) Summary of the top HOMER known motifs enriched in ESR1 peaks in THESC^ESR1^ treated with E2 for 3 hours. (G) Venn diagram comparing all genes whose TSS is within 100 kb of ESR1 peaks in THESC^ESR1^ (treated with E2 for 3 hours) and genes regulated by ESR1/E2 in THESC^ESR1^ (E2-dependent DEGs). THESC = immortalized human endometrial stromal cells; THESC^ESR1^ = ESR1 activated cells; E2 = 10 nM estradiol; vehicle = 0.1% ethanol; TSS = transcription start site.

To identify genes regulated by ESR1/E2 with nearby ESR1 binding, ESR1 peaks were annotated to all genes whose TSS was within 100kb and overlapped with E2-dependent DEGs. 134 DEGs had at least one ESR1 binding site within 100kb of their TSS (Figure 3G). These genes were enriched for pathways including ERa genomic signaling, inflammatory response, vasculature development, the core matrisome, and homeostasis (Table 2). These results support ESR1/E2’s role as a key regulator of stromal cell homeostasis and preparation for pregnancy.

**Table 1.2:**
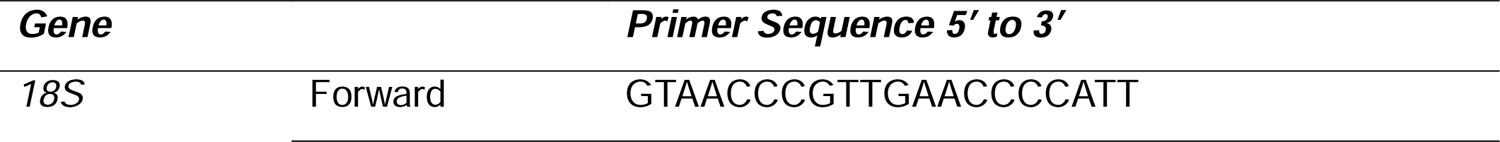
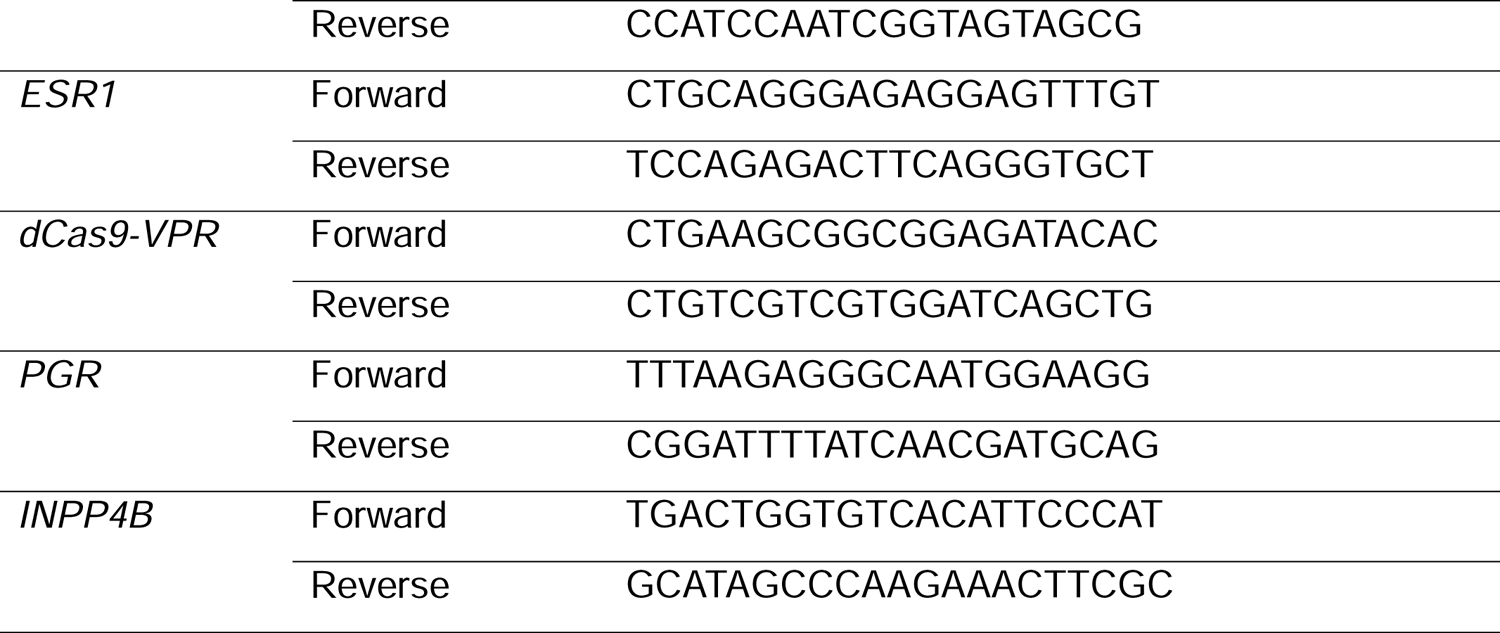
Primer sequences for RT-qPCR.

### Chromatin landscape in hESC and decidualized hESC and association with ESR1 transcriptome and cistrome in engineered THESC

Chromatin conformation capture-based methods such as Hi-C and HiChIP have emerged as useful tools to identify distal genome contact points that form promoter-enhancer interactions and topologically associated domains (TADs) (57). To investigate the 3D chromatin landscape in hESC, we performed H3K27ac HiChIP in two primary hESC cell-lines treated with vehicle (0.02% EtOH) or a decidualization cocktail (10 nM E2, 1 uM MPA, 100 uM cAMP; EPC) for 72 hours. We identified 150,966 and 193,367 loops in vehicle- and EPC-treated cells, respectively (Figure S4; Dataset 5). Differential analysis (|FC| > 2 and p-value < 0.05) identified 1,421 EPC-repressed and 2,107 EPC-activated loops (Figure 4A; Dataset 5). We focused on the differential loops that had at least one anchor at a gene’s promoter (26% of EPC-repressed and 14% of EPC-activated loops; Figure 4B) and overlapped these genes with DEGs in response to EPC treatment for 72 hours from a published RNAseq dataset (GSE205481) in two matched donor hESC, and a third replicate (Dataset 6) (26). There were 141 DEGs in response to EPC treatment that also had a differential promoter-contacting loop (Figure 4C). Most EPC-activated DEGs were linked to EPC-activated loops (98%), and vice versa for repressed DEGs (93%) (Figure 4D-E). IPA pathway analysis highlighted enrichment in immune signaling, prolactin signaling, and endometrial remodeling pathways including vasculature development and integrin interactions (Figure 4F). These findings suggest that chromatin architecture reorganization supports gene expression changes driving decidualization and endometrial remodeling.

**Figure 4.**
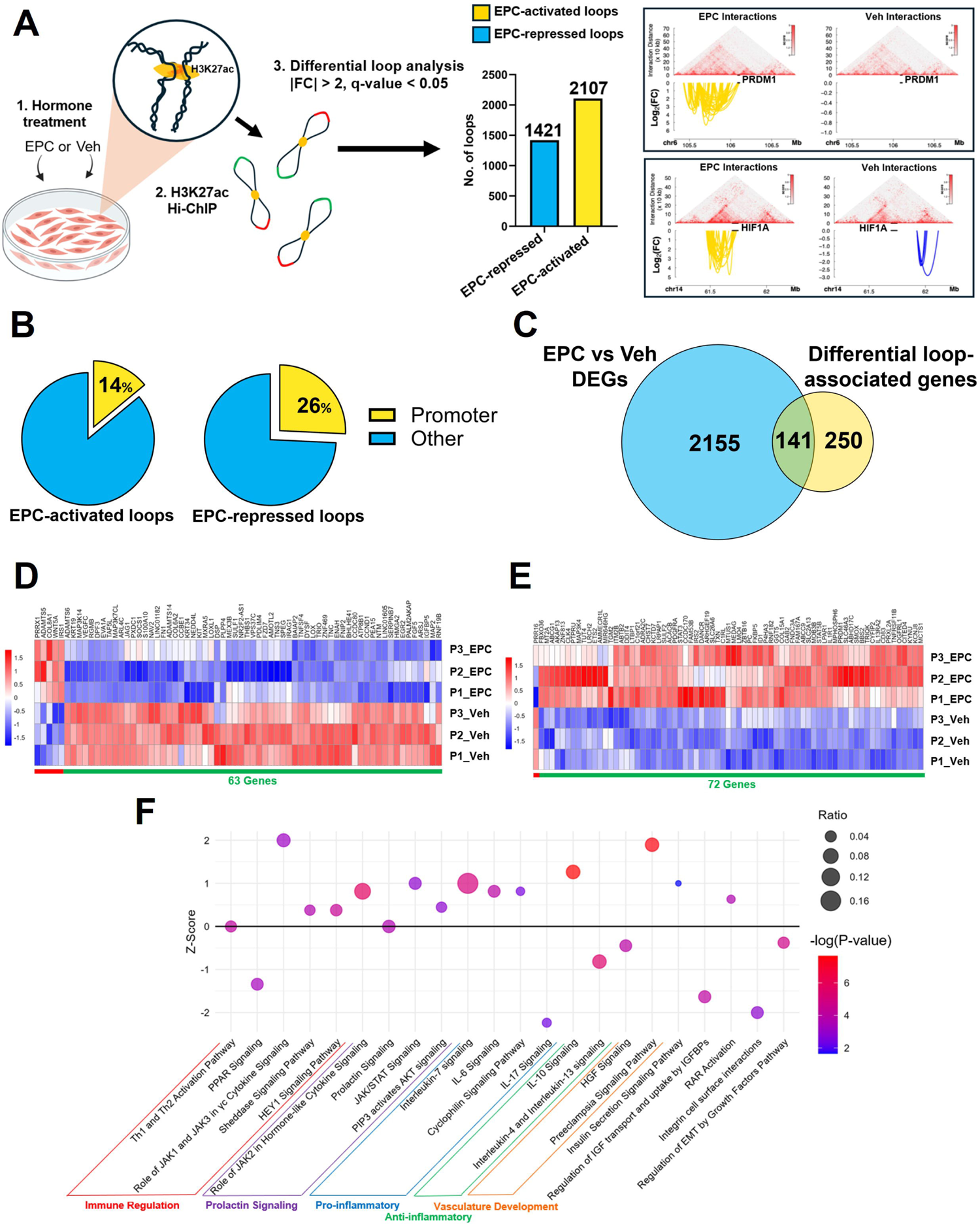
Chromatin landscape in hESC and decidualized hESC assessed through H3K27ac HiChIP. (A) H3K27ac HiChIP protocol in hESC treated with Veh or EPC. Raw contact matrices for Veh or EPC samples (n=2 merged independent samples per matrix) and significant differential loop interactions around genes of interest. Labeled black lines indicate genes of interest from transcription start site to transcription end site. (B) Pie charts showing the percentage of loops with at least one anchor at the promoter of a gene (orange) in EPC-activated and repressed loops. (C) Comparison between genes with a differential promoter-targeting loop in hESC and genes differentially expressed in response to EPC from previously published RNA-seq (GSE205481). (D) Heatmap generated via unsupervised clustering of genes differentially expressed in response to EPC that contain an EPC-repressed promoter targeting loop or (E) EPC-activated promoter targeting loop. (F) Select pathways from IPA pathway analysis of 141 genes differentially expressed in response to EPC with a differential promoter-targeting loop. EPC = 10 nM E2, 1 uM medroxyprogesterone acetate, and 100 uM cAMP; IPA = ingenuity pathway analysis.

Estrogen signaling has been shown to be critical for decidualization. To identify genes regulated by E2 that may be involved in the regulation of decidualization, we integrated the EPC H3K27ac HiChIP loops, ESR1 binding sites, and ESR1/E2 transcriptome to identify ESR1/E2 target genes. We then overlapped these genes with genes that are differentially expressed in response to EPC. To identify ESR1/E2 target genes, we focused on genes that have an ESR1 binding peak at the promoter of the gene (proximal binding; 89 genes), or an ESR1 binding peak that is one HiChIP loop away from the promoter (anchored binding; 1,030 genes) (Figure 5A; Dataset 5). We then overlapped proximal and anchored genes with the 369 genes regulated by ESR1/E2 in the engineered THESC (ESR1-E2 vs ESR1-Veh). We identified 72 genes regulated by ESR1/E2, with 86% of these genes activated in response to E2 (Figure 5A-B). To identify if any of these genes are involved in the process of decidualization, we compared them to genes that are differentially expressed in response to EPC. 28 genes overlapped with DEGs in response to EPC (Figure 5C). We identified genes involved in endometrial remodeling and decidualization, including FOXO1 and IL6R that are upregulated in response to E2 and EPC, and have proximal or anchored ESR1 binding sites at the promoter (Figure 5D-E). These results suggest that distal ESR1 binding sites may regulate gene expression for decidualization through long-range chromatin loops.

**Figure 5:**
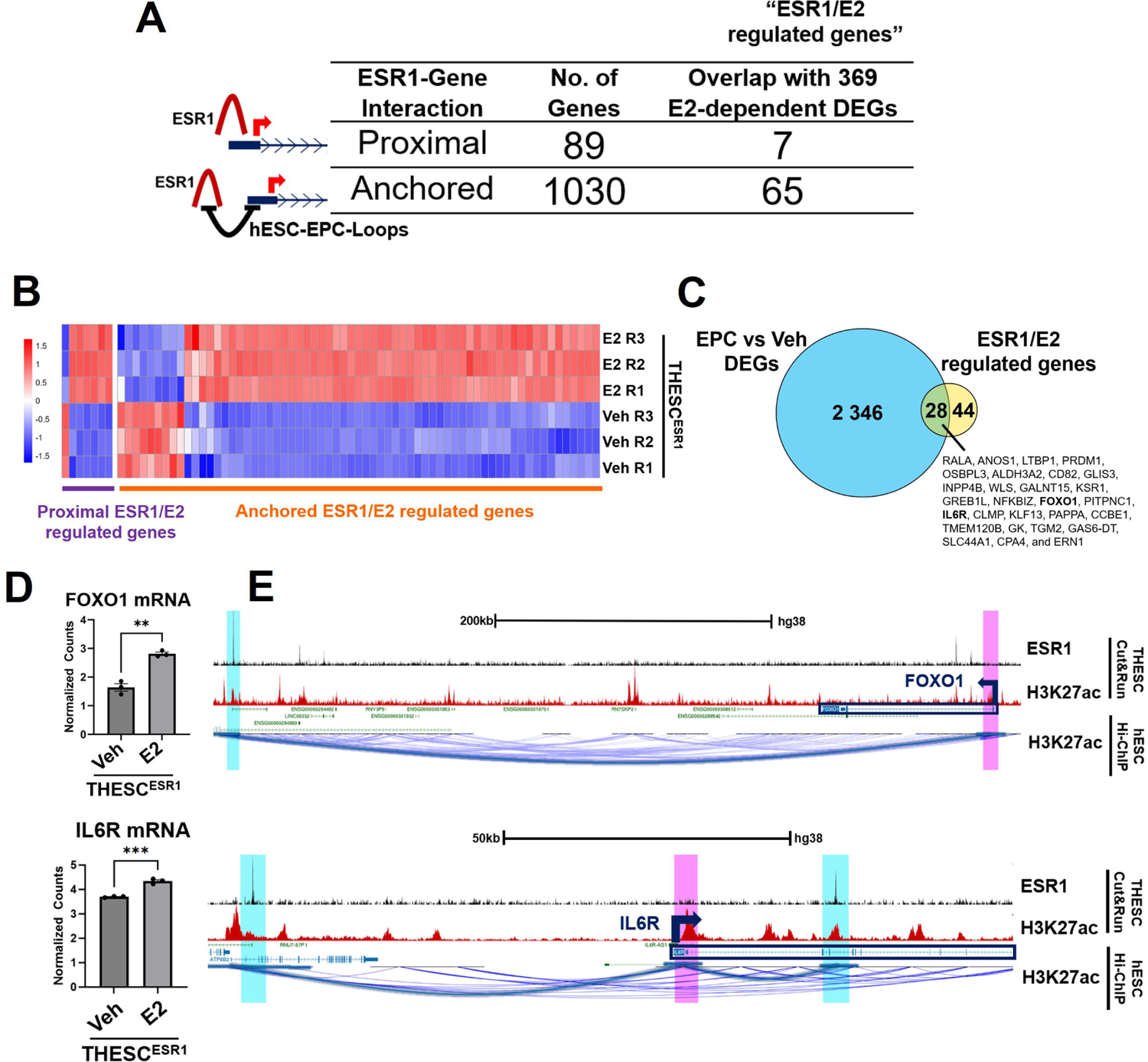
Integration of ESR1 Cut&Run, RNA-seq, and H3K27ac EPC HiChIP loops to identify genes regulated by ESR1/E2 involved in decidualization. (A) Table showing the number of genes with proximal or anchored ESR1 binding (identified through integration of H3K27ac HiChIP and ESR1 Cut&Run), as well as overlap with 369 genes regulated by ESR1/E2 in THESC^ESR1^ (E2-dependent DEGs). (B) Heatmap generated via unsupervised hierarchical clustering of 80 genes regulated by ESR1/E2 with proximal or anchored ESR1 binding at the promoter. (C) Comparison between genes regulated by ESR1/E2 with proximal or anchored ESR1 binding at the promoter, and genes differentially expressed in response to EPC from previously published RNA-seq (GSE205481). (D) Bar chart comparing RNA-seq log2CPM normalized counts for FOXO1 and IL6R in THESC^ESR1^ treated with E2 or vehicle for 24 hours. Error bars are shown as mean with SEM. (E) UCSC genome browser views around the FOXO1 and IL6R loci. Tracks show ESR1 Cut&Run in THESC^ESR1^ treated with E2 for 3 hours, with IgG background subtracted (black), H3K27ac Cut&Run in THESC (red), and H3K27ac HiChIP loops in EPC treated hESC (blue). The promoter of the gene of interest is highlighted in pink, ESR1 binding peaks are highlighted in blue, and the HiChIP loop that connects the gene promoter with the ESR1 binding peak is highlighted with a blue loop. THESC = immortalized human endometrial stromal cells; EPC = 10 nM E2, 1 uM medroxyprogesterone acetate, and 100 uM cAMP; THESC^ESR1^ = ESR1 activated cells; E2 = 10 nM estradiol; CPM = counts per million; SEM = standard error of the mean.

To investigate ESR1/E2’s regulation of genes in the endometrial stroma beyond decidualization, we identified ESR1/E2 target genes using the same approach as described, with a focus on HiChIP loops in vehicle-treated hESC. We identified 69 genes that are differentially expressed in response to E2 in THESC with proximal or anchored ESR1 binding at the promoter (Figure 6A). Metascape pathway analysis showed an enrichment for cancer pathways including P53 signaling, endometrial neoplasms, and breast cancer (Figure 6B). We identified genes that are implicated in endometrial cancer including ERRFI1 and NRIP1, as well as genes involved in other types of metastatic cancers, such as EPAS1, that are regulated by ESR1/E2 (Figure 6C-D). Lastly, we show that ESR1, both in the presence and absence of E2, increases cell viability in THESC through MTT assay (Figure 6E). We also demonstrate that in response to E2, ESR1 significantly enhances the migration of THESC through a porous membrane, as assessed by a transwell migration assay (Figure 6F).

**Figure 6:**
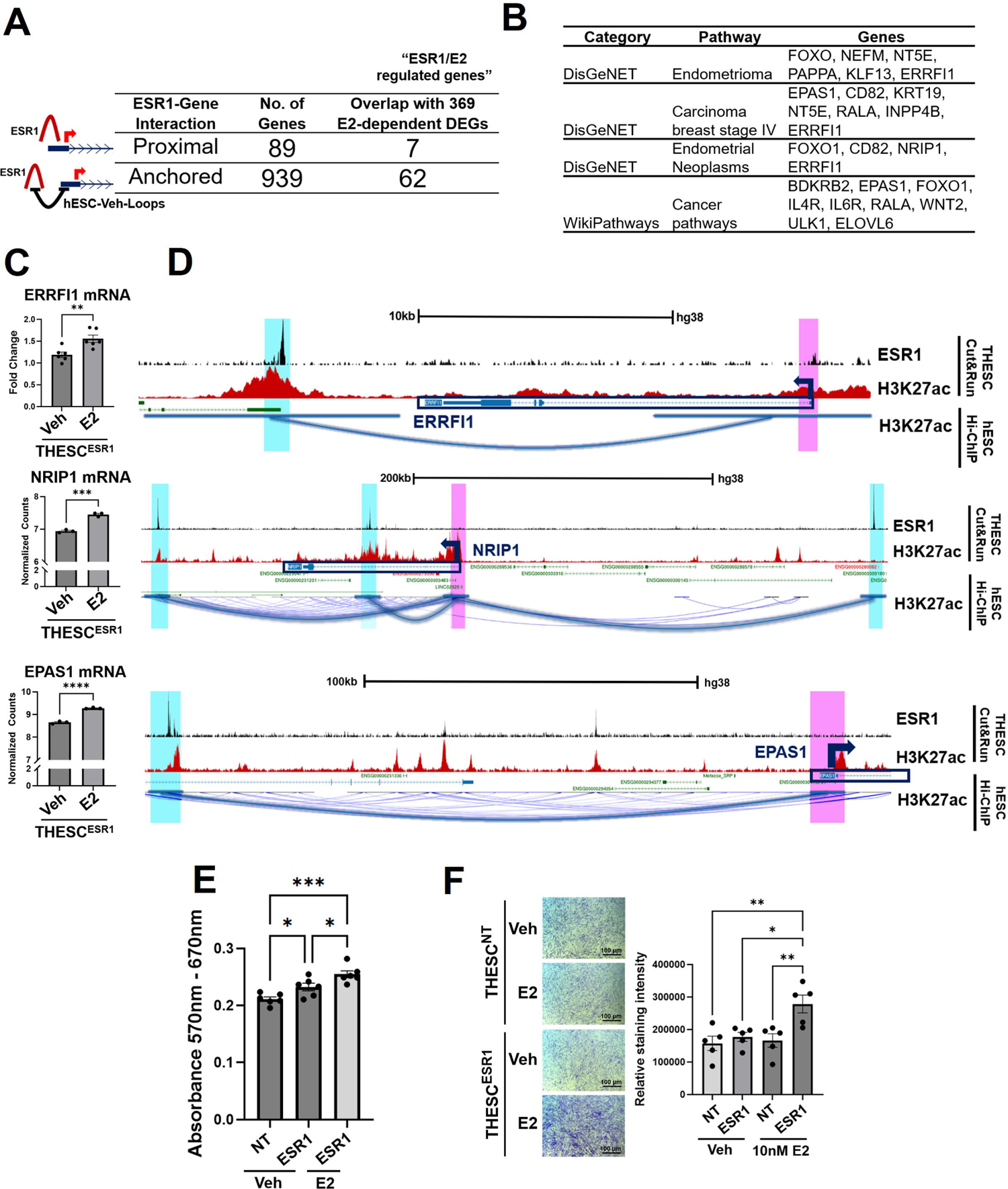
Integration of ESR1 Cut&Run, RNA-seq, and H3K27ac HiChIP loops to identify genes regulated by ESR1/E2 that are implicated in endometrial cancer. (A) Table showing the number of genes with proximal or anchored ESR1 binding (identified through integration of H3K27ac vehicle HiChIP and ESR1 Cut&Run), as well as overlap with 369 genes regulated by ESR1/E2 in engineered THESC (E2-dependent DEGs). (B) Select pathways from metascape pathway analysis of 75 genes regulated by ESR1/E2 with proximal or anchored ESR1 binding at the promoter. (C) Bar chart comparing RNA-seq log2CPM normalized counts for ERRFI1, NRIP1, and EPAS1 in THESC^ESR1^ treated with E2 or vehicle for 24 hours. Error bars are shown as mean with SEM. (D) UCSC genome browser views around the ERRFI1, NRIP1, and EPAS1 loci. Tracks show ESR1 Cut&Run in THESC^ESR1^ treated with E2 for 3 hours, with IgG background subtracted (black), H3K27ac Cut&Run in THESC (red), and H3K27ac HiChIP loops in vehicle treated hESC (blue). The promoter of the gene of interest is highlighted in pink, ESR1 binding peaks are highlighted in blue, and the HiChIP loop that connects the gene promoter with the ESR1 binding peak is highlighted with a blue loop. (E) Cell viability assessed by MTT assay in THESC^ESR1^ and THESC^NT^ treated with vehicle or E2 for 4 days. Statistical significance was determined by one-way ANOVA with significance defined as p-value < 0.05. Error bars are shown as mean with SEM. (F) Microscopic images and quantification of density of migrated THESC^ESR1^ and THESC^NT^ treated with vehicle or E2 for 72 hours. Cells were stained with 1% crystal violet solution, and cell density was quantified using ImageJ. Statistical significance was determined by one-way ANOVA with significance defined as p-value < 0.05. Error bars are shown as mean with SEM. THESC = immortalized human endometrial stromal cells; THESC^ESR1^ = ESR1 activated cells; SEM = standard error of the mean; THESC^NT^ = control non-targeting cells; E2 = 10 nM estradiol; vehicle = 0.1% ethanol; EPC = 10 nM E2, 1 uM medroxyprogesterone acetate, and 100 uM cAMP; CPM = counts per million; SEM = standard error of the mean.

These findings highlight a key role for ESR1/E2 signaling in regulating gene expression in the endometrial stroma during decidualization and in disease. The identification of ESR1/E2 target genes enriched in cancer-related pathways suggests estrogen signaling in the stroma might be associated with disease progression, potentially through an increase in stromal proliferation and migration.

## DISCUSSION

Estrogen signaling through its receptor ESR1 is critical for successful pregnancy by driving decidualization, implantation, and uterine receptivity in the endometrium (46). Low estrogen levels are suspected to contribute to abnormal placentation in naturally conceived pregnancies, whereas an excess of estrogen may impair pregnancy development and lead to adverse outcomes (17). Moreover, its dysregulation is linked to gynecological pathologies, including endometriosis and endometrial cancer (47, 48).

Studying E2 signaling in endometrial stromal cells has been hindered by the limited E2 responsiveness of both primary and immortalized cell lines due to the silencing of ESR1 in vitro (19). In this study, we provide novel insights into ESR1-mediated transcriptional regulation through RNA-seq and Cut&Run assays in human endometrial stromal cells (hESCs) with restored ESR1 expression and E2 responsiveness using CRISPR. We also profiled the chromatin landscape in hESC and decidualized hESC through H3K27ac HiChIP. We integrated this data with the ESR1 cistrome to associate distal ESR1 binding peaks with the promoters of the genes they regulate through long-range looping. We identified genes critical for decidualization and implicated in endometrial cancer that are regulated by ESR1/E2 with proximal or anchored ESR1 binding up to 500 kb away from their promoter.

Over the past decades, studies in breast cancer cells have significantly expanded ESR1/E2 actions beyond the classical model of ESR1 transcriptional regulation, which originally involved ligand activation and binding to EREs (49), to include ligand-independent actions and binding to non-ERE genomic sites through protein-protein interactions (50). Through RNA-seq and Cut&Run assays, we show that ESR1 can bind to the genome and regulate transcription in both the presence and absence of ligand in THESC. Importantly, comparison of DEGs between control and E2-treated conditions in ESR1-activated hESCs revealed that 72% of genes overlapped with those active in human endometrial tissue during the proliferative phase, emphasizing the physiological relevance of our model. Using HOMER, we identified transcription factor motifs enriched in ESR1 binding sites in THESC, including chromatin modifiers and proteins known to regulate ESR1 tethering through protein-protein interactions, such as HOX, JUN/AP-1, RUNX, and TEAD motifs (42–44). Additionally, we found enrichment for zinc-finger GATA transcription factors that are implicated in endometrial function and pathology (51). These results demonstrate that ESR1 activation restores E2 responsiveness in THESC, establishing this model as a valuable tool for studying estrogen signaling in endometrial stromal cell biology.

Through H3K27ac HiChIP, we also show that changes to chromatin architecture support gene expression changes that drive decidualization. Integrating differential chromatin loops with DEGs in response to EPC from published RNA-seq (GSE205481) showed enrichment in inflammatory and immune pathways, including prolactin and its downstream signaling pathway (JAK/STAT, PIP3/AKT), as well as pro-inflammatory interleukin signaling including IL-6 and IL-7 signaling (26). Blastocyst implantation, decidualization, and early pregnancy relies on a pro-inflammatory state mediated in part by cytokine release from immune cells (58). Prolactin signaling is a key marker of decidualization and it’s release during the secretory phase coincides with the onset of endometrial decidualization (59). Prolactin and its receptor are indispensable for pregnancy (60) and play roles in regulating angiogenesis, glandular secretion, and immune regulation (61). Taken together, our results suggest that changes to the chromatin landscape may be involved in regulating inflammatory pathways for early pregnancy establishment and decidualization.

It has become common practice to associate transcription factor binding peaks to nearby genes, but this approach can introduce biases, particularly for regulators like ESR1 that bind to distal cis-regulatory elements. To avoid assumptions introduced through arbitrary association of binding peaks with nearby genes, we integrated H3K27ac HiChIP with ESR1 Cut&Run data in engineered THESC. We show that chromatin looping may be involved in bringing distal ESR1 peaks to promoters of genes transcriptionally regulated by ESR1/E2. One notable example is FOXO1; FOXO1 is activated in response to ESR1/E2 signaling and HiChIP data suggest that ESR1 regulation of FOXO1 may occur through a long-range loop connecting the FOXO1 promoter to an ESR1 binding site located more than 500kb away from the FOXO1 TSS. We have previously shown that FOXO1 is a critical co-regulator of PGR during decidualization (62) and it is involved in the activation of decidual markers PRL and IGFBP1 (63–66). FOXO1 is also involved in PGR’s anti-proliferative effects on the endometrial stroma and epithelium, and its misregulation is implicated in endometrial cancer (67, 68). FOXO1 expression can be induced by 8-Br-cAMP (63), BMP4 (69), and SOX4 (70) in the endometrium, and here we suggest that liganded ESR1 is also a regulator of FOXO1 in endometrial stromal cells. Taken together, these results suggest that changes to chromatin architecture may be involved in the regulation of genes for endometrial differentiation. Furthermore, we demonstrate that the practice of integrating cistromic data with chromatin looping may be a less arbitrary method to associate distal transcription factor binding sites with the promoters of genes they regulate.

Through the integration of HiChIP, ESR1 Cut&Run, and RNAseq data, we also identified genes regulated by ESR1/E2 that are implicated in endometrial cancer, including genes NRIP1 (RIP140) and ERRIF1 (MIG-6). We found three distal ESR1/E2 binding sites located approximately 60kb, 200kb, and 270kb away from the NRIP1 TSS that are brough to the NRIP1 promoter through HiChIP chromatin looping. NRIP1 functions as a repressor of ESR1 signaling to attenuate its expression (71, 72) and has high rates of mutation in tumors (73). NRIP1 silencing is implicated in the progression of endometrial cancer (73), although it is suggested to act as both an activator and inhibitor of tumorigenesis in various cancers (74–76). In the case of ERRFI1, we identified an ESR1/E2 binding site approximately 20kb away from the ERRFI1 TSS with HiChIP chromatin looping. ERRFI1 mediates PGR’s repression on E2-induced proliferation in the endometrium and its knockdown in the uterus of mice leads to endometrial hyperplasia (77). In humans, ERRFI1 downregulation is correlated with human breast carcinoma (78, 79), complex atypical hyperplasia, and endometrioid endometrial carcinomas (77). ESR1/E2’s regulation of ERRFI1 has previously been documented in the chicken oviduct (80). Our study suggests that ESR1/E2 regulates the genes NRIP1 and ERRFI1, both of which are involved its negative feedback mechanism. Disruption of ESR1/E2 regulation of these genes may occur in endometrial cancer, resulting in unopposed estrogen signaling that drives tumorigenesis. Although endometrial cancer originates from the epithelium, the growth of the epithelium strongly depends on paracrine signaling from the endometrial stroma. The genes identified may play a role in early signaling events during endometrial cancer development and could contribute to epithelial mis regulation that drives tumorigenesis.

## Conclusion

In this study, we developed an in vitro model by engineering THESC expressing the CRISPR activation system to activate ESR1, restoring their responsiveness to E2. Integration of ESR1/E2 transcriptome and cistrome with HiChIP data identifies its role in regulating inflammation, proliferation, and decidualization, as well as its implications in endometrial cancer, providing new insights into estrogen-mediated transcriptional regulation and chromatin organization. This model serves as a powerful tool to study estrogen signaling in endometrial stromal biology and related pathologies.

## ACKNOWLEDGMENTS

This work was supported by an Intramural Research Program of the National Institute of Environmental Health Sciences (NIEHS), National Institutes of Health (NIH) 1Z1AES103311 (FJD). We thank the following NIEHS core facilities for their exceptional technical support: the Epigenomics and DNA Sequencing Core Facility, supported by 1ZICES102545-17, the Integrative Bioinformatics Support Group, supported by 1ZICES103371, and the Viral Vector Core, supported by ZIC ES102506-09. We also thank Elvis Quiroz for technical assistance.

---

**Figure S1:** Full length uncropped western blot.

**Figure S2.** RT-qPCR analysis of ESR1 expression in engineered immortalized human endometrial stromal cells (H1644) and myometrial cells (hTERT-HM).

**Figure S3.** Unsupervised hierarchical clustering and multidimensional scaling plot of RNAseq data.

**Figure S4.** Raw and filtered contact matrices comparing genome-wide interactions in Veh and EPC treated hESC.

**Dataset S1:** (A) DEGs (Adjusted P-value <0.05, |Fold-Change| > 1.3) between ESR activated cells treated with Vehicle for 24 hours versus control cells treated with Vehicle for 24 hours. (B) IPA pathway analysis of DEGs.

**Dataset S2:** (A) DEGs (Adjusted P-value <0.05, |Fold-Change| > 1.3) between ESR activated cells treated with 10nM Estradiol for 24 hours versus Vehicle for 24 hours. (B) IPA pathway analysis of DEGs.

**Dataset S3**: 220 DEGs regulated by E2-independent ESR1 and 268 DEGs regulated by E2-dependent ESR1 in THESC, that are also active (FPKM > 1) in human endometrial tissue during the proliferative stage (GSE132713).

**Dataset S4:** ESR1 ESR1 Cut&Run peaks in THESC engineered to express ESR1 and treated with E2 or vehicle (0.01% EtOH) for 1, 3, or 6 hours. Columns A-C indicates peaks in bed format. Column D (tssDistance) indicates how far the peak is from the nearest transcription start site. Column E (relativeLocation) indicates the genomic feature that the peak overlaps with. Column F (overlappedPromoters) indicates the peak a gene overlaps with if it is located at the gene promoter. Column G (closestTSS) indicates the single closest gene transcription start site. Column H (genesWithTSSWithin100kb) annotates each peak to all the genes with a transcription start site that are within 100 kb of the peak. Columns I and K (Veh and EPC_annotated_loopedPromoterGenes) indicate whether the peak overlaps with an H3K27ac Hi-ChIP loop anchor that anchors to a gene promoter in (in Veh or EPC H3K27ac HiChIP). Columns J and L (Veh and EPC_annotated_loopIDs) indicates which HiChIP loop brings the peak to the gene promoter.

**Dataset S5**: H3K27ac Hi-ChIP Loops in hESC treated with vehicle (0.02% EtOH) or EPC, as well as the loops that are significantly activated or repressed in response to EPC treatment in comparison to vehicle. Columns A - F indicate the loop anchors for each loop in bed format. Column G (loopID) assigns a unique ID to each loop. Columns H and I (overlappedGenes1/2) If the loop anchor overlaps with any part of a gene, the gene is indicated here. Columns E and J (relativeLocation1 and 2) indicates the genomic feature that each loop anchor overlaps with.

**Dataset S6:** Reanalysis of published RNAseq (GSE205481) comparing gene expression in hESC treated with EPC versus vehicle (GSM6213359, GSM6213360, GSM6213363, GSM6213364, GSM6213367, GSM6213368).

