## Supplementary figures and images for "Deciphering ESR1-driven transcription in human endometrial stromal cells via transcriptome, cistrome, and integration with chromatin landscape"

### Figure_S1

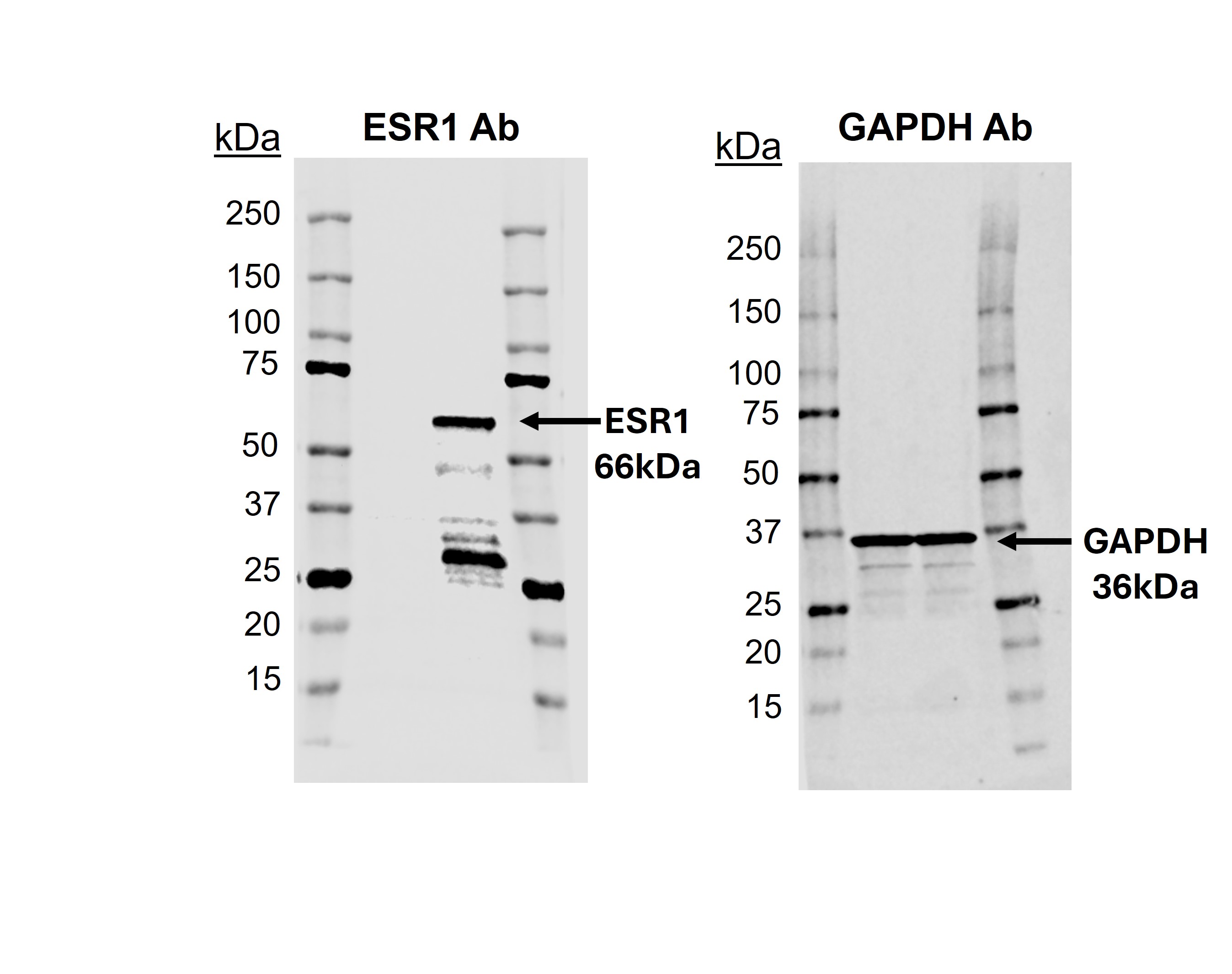

### Figure_S2

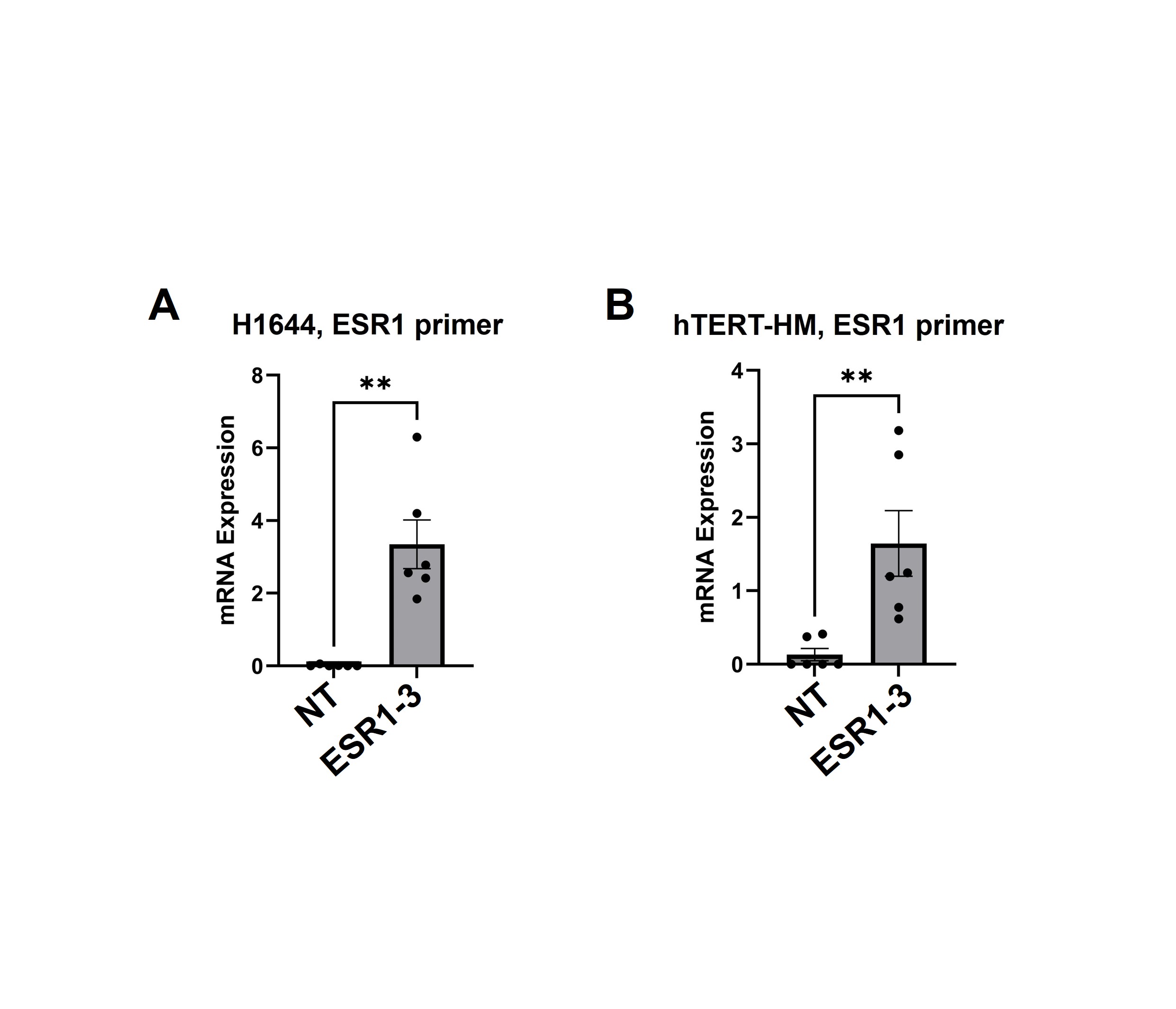

### Figure_S3

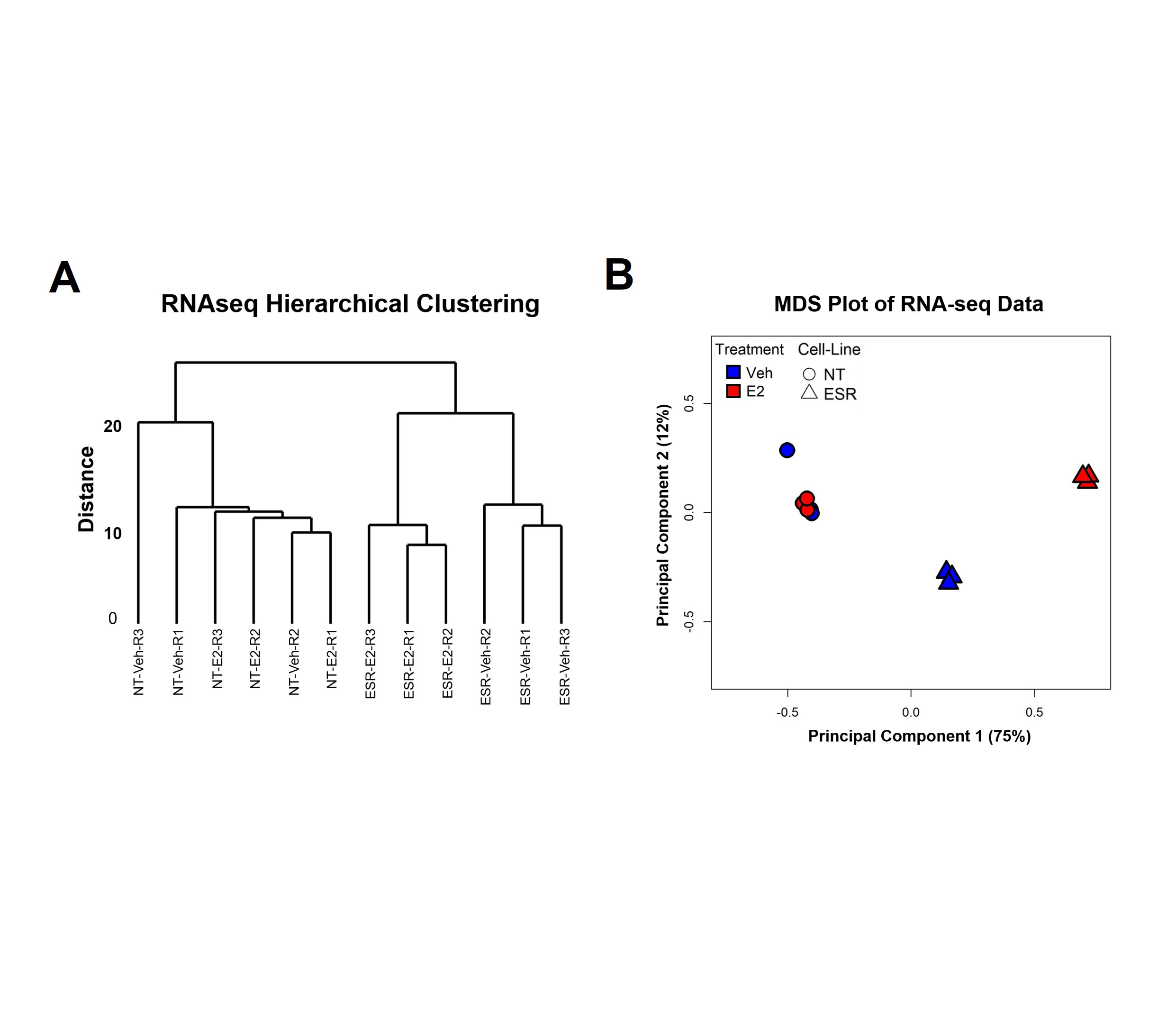

### Figure_S4

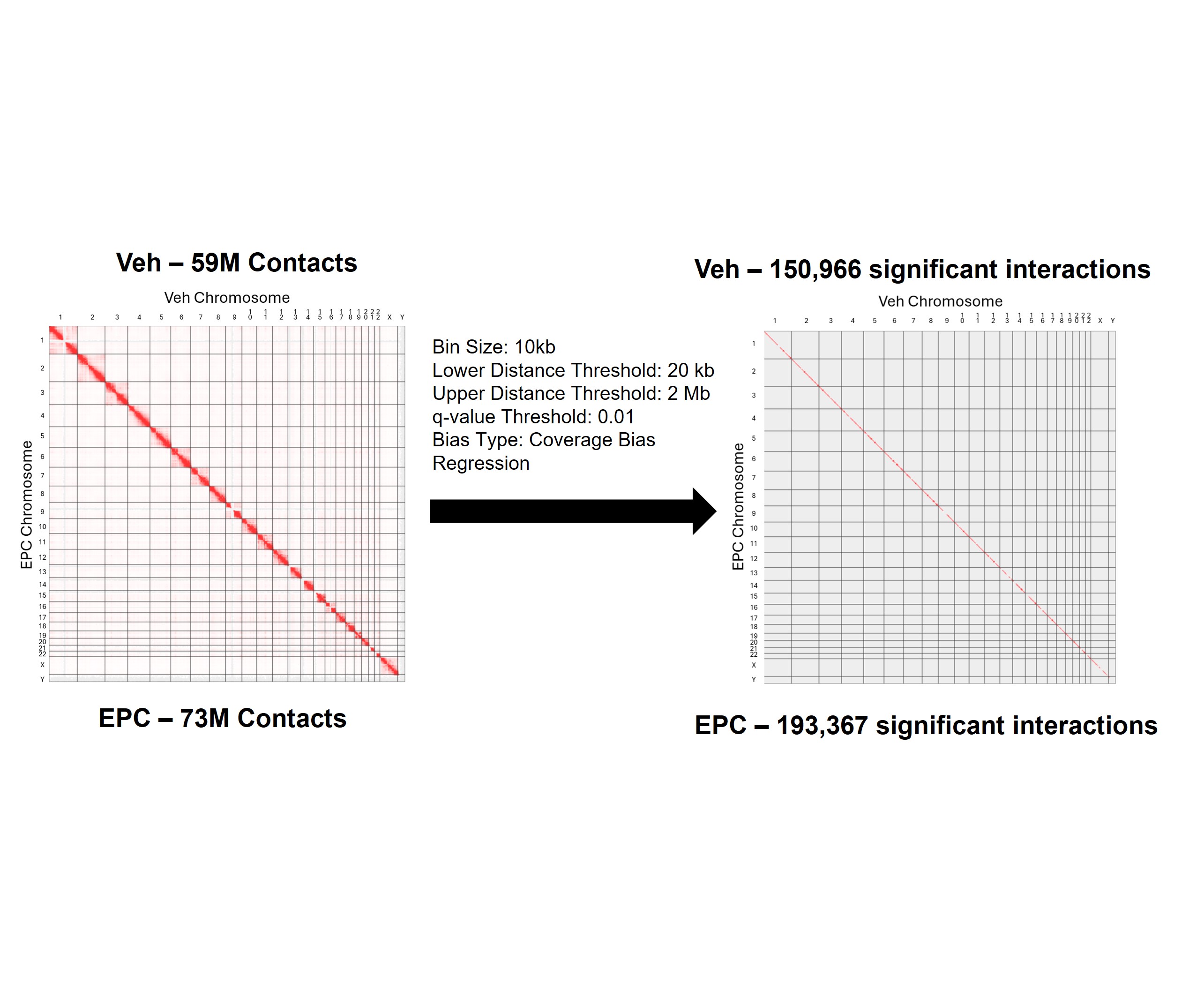
